# Representation of music genres based on the spectro-temporal modulation responses of the human brain

**DOI:** 10.1101/471326

**Authors:** Tomoya Nakai, Naoko Koide-Majima, Shinji Nishimoto

## Abstract

Music genre is an essential category for understanding human musical preferences and is provided based on the abstract categorization upon complex auditory stimuli. Previous neuroimaging studies have reported the involvement of the superior temporal gyrus (STG) in response to general music-related features. However, it remains largely unclear how abstract categories of music genre are represented in the brain and what acoustic features are more suited for explaining such representations. Here we examined comprehensive cortical representations and functional organization of music genres using 540 music clips. We applied a voxel-wise modeling approach to music-evoked brain activity measured using functional magnetic resonance imaging (fMRI). We observed distinct cortical organizations for different music genres in the bilateral STG, which revealed the representational relationship between various music genres, e.g., classical and hip-hop music showed opposite representations. Representations of music genres were largely explained by spectro-temporal modulation, which was modeled by a biologically plausible spectro-temporal modulation-transfer function (MTF) model. Our results elucidate the quantitative representation of music genres in the human cortex and indicate the possibility of modeling our abstract categorization of complex auditory stimuli based on the brain activity.

**Significance statement:** Music genre is an essential category for understanding human preferences of music. However, it is largely unknown how abstract categories of music genre are represented in the brain. Here, we examined comprehensive cortical representations of music genres by building voxel-wise models of fMRI data collected while human subjects listened to 540 music clips. We found distinct cortical organizations for various music genres in the bilateral STG. Such genre-specific cortical organization was explained by the biologically plausible MTF model. The current study elucidates the quantitative representation of music genres in the human cortex for the first time and indicates the possibility of modeling our abstract categorization of complex auditory stimuli based on the brain activity.

## Introduction

When we listen to music, we tend to classify them into genre categories like classical, rock, or hip-hop. Thus, music genre is an essential class-label for understanding human preferences of music, and it has been widely used in research on music information retrieval (1). However, how such abstract genre categories were perceived and how our brain subserves this classification process remain largely unknown.

Previous neuroimaging studies have investigated brain representations of general music-related features such as loudness, timbre, harmony, and rhythm (2–4). For example, Alluri et al. (2012) have reported that activation in the bilateral superior temporal gyrus (STG) was significantly correlated with timbre, harmony, and rhythm features (2). Moreover, Toiviainen et al. (2014) have demonstrated that the bilateral STG contributed to the decoding of timbre features (3). Together, these studies have indicated the involvement of the bilateral STG in response to such music-related features.

In the physiology-oriented literature studying acoustic representation of STG, the spectro-temporal modulation-transfer function (MTF) model has been widely applied as a biologically plausible model (5–8). Modulation-selective responses have been observed in the primary auditory cortex of ferrets (9) and in the human brain, using both ECOG (10, 11) and fMRI (12, 13). The MTF model has also been used to explain brain activation differences between 2-s excerpts of music and voices (7) as well as those between simple tones of different musical instruments (5). However, it is not yet certain whether the MTF model is more suited for explaining brain activity patterns of abstract genre categories, which are composed of complex auditory stimuli.

Recent neuroimaging studies have used voxel-wise encoding/decoding-models (14) to examine sensory and higher-order cortical representations, including visual (15, 16) and auditory modalities (17, 18). An encoding/decoding model approach has the advantage that it can compare the performances of several competing theoretical models using the same dataset. For example, de Heer et al. (2017) modeled brain activation during passive story listening and performed encoding model fitting using cochlear, phoneme, and semantic models (17).

Through such an approach, we can evaluate whether a biologically plausible model such as MTF is more effective than other models for predicting music genre-representing brain activation.

In the present study, we applied the encoding/decoding model approach to study brain activity induced by music stimuli of various genres and then examined the detailed cortical organization underlying each genre. Five subjects passively listened to naturalistic music stimuli representing 10 different music genres, and brain activation was quantified using fMRI (Fig. 1A). We hypothesized that pieces of music are represented in the human brain in a genre-specific way and recognized based on their spectro-temporal modulation. We first examined specific cortical activation patterns based on predefined genre-labels (Fig. 1B). Acoustic features were then extracted using two biologically plausible models (cochlear, MTF) and two music-related models [music information retrieval (MIR), mel-frequency cepstral coefficient (MFCC)]. We then investigated whether the MTF model most accurately explained the genre-specific cortical organization. We further examined the representational specificity among music genres by performing genre classification with brain activity, behavior, and extracted features. Finally, we tested whether such representational differences of music genres were also captured by spectro-temporal modulation. Preliminary results of the current study were *in press* in IEEE International Conference on Systems, Man, and Cybernetics (IEEE SMC 2018) (19).

**Fig. 1.**
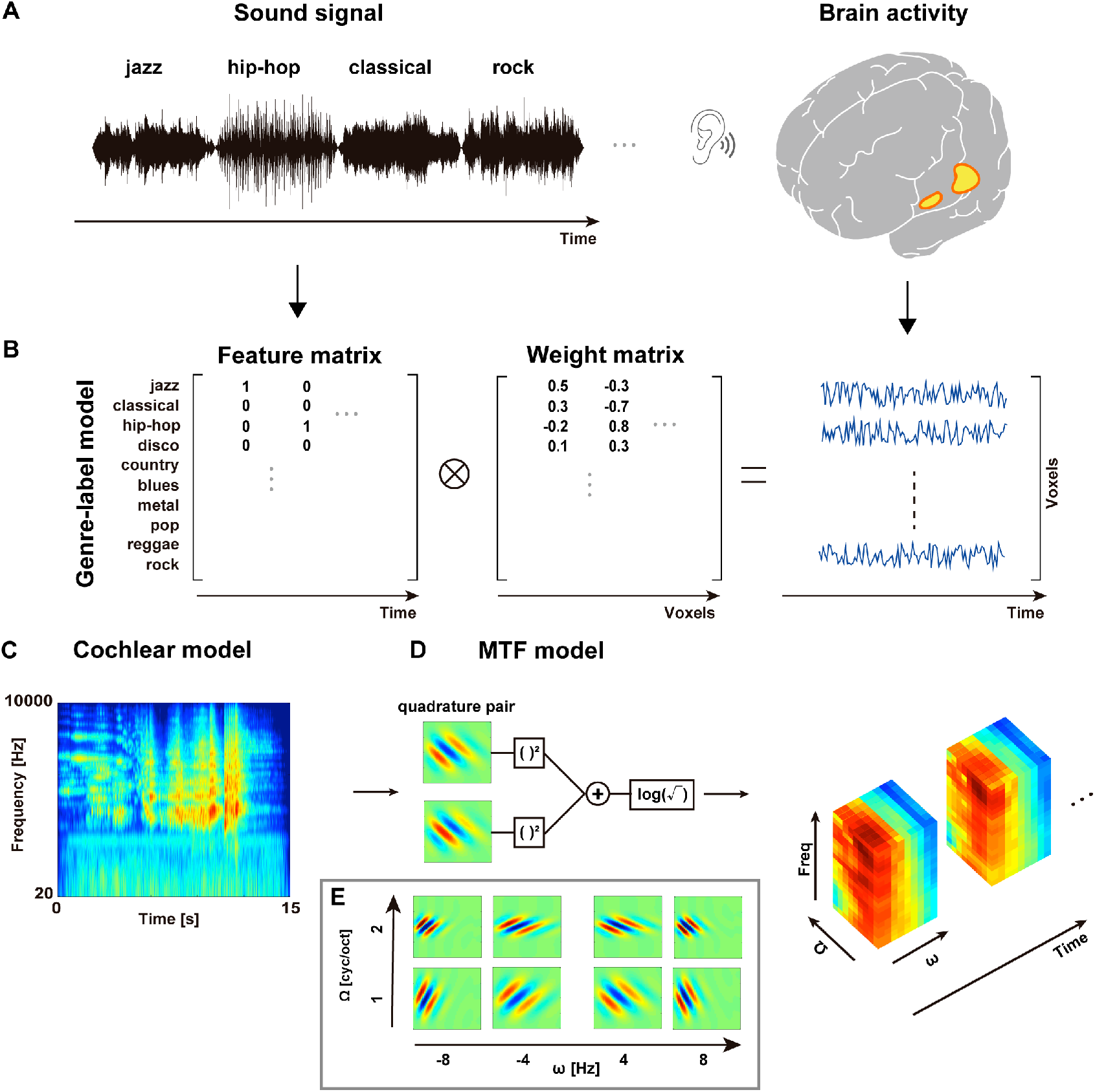
Schematic image showing the research paradigm of the current study. **(A)** Subjects passively listened to the naturalistic music stimuli of 10 music genres and the evoked brain activity was measured using fMRI. **(B)** Voxel-wise brain activity was modeled as a feature matrix (music genre-labels) times a weight matrix. Regularized linear regression was used to estimate the optimal weights. **(C)** An example cochleogram, which was used as the acoustic feature in the cochlear model. **(D)** modulation-transfer function (MTF) model features were extracted from each cochleogram. Modulation energy was calculated by squaring and summing the outputs of modulation-selective filters in quadrature. Modulation energy was then passed through a compressive nonlinearity, averaged across 1.5 s, and further averaged within each of the 20 non-overlapping frequency ranges logarithmically spaced along the frequency axis, resulting in a series of 2000 dimensional MTFs. **e**, Examples of the modulation-selective filters with four temporal modulation parameters (ω (Hz)) and two temporal modulation parameters (Ω (cyc/oct)). Note that the upward and downward (i.e., with positive and negative ω, respectively) filter outputs were averaged to calculate MTF.

## Results

### Genre-representing cortical organization

Genre-representing cortical areas were assessed using the genre-label model. In all subjects, significant prediction accuracy was observed in the bilateral STG (*p* < 0.05, with FDR correction; Fig. 2A and Supplementary Fig. 1). To identify cortical areas robustly representing music genres independent of sample selection, we determined the genre-representing functional ROI for each subject (Fig. 2B and Supplementary Fig. 2) using a resampling procedure (see Methods). This analysis again demonstrated significant prediction accuracies in the bilateral STG. The functional ROI was used as an inclusive mask in the subsequent analyses.

**Fig. 2.**
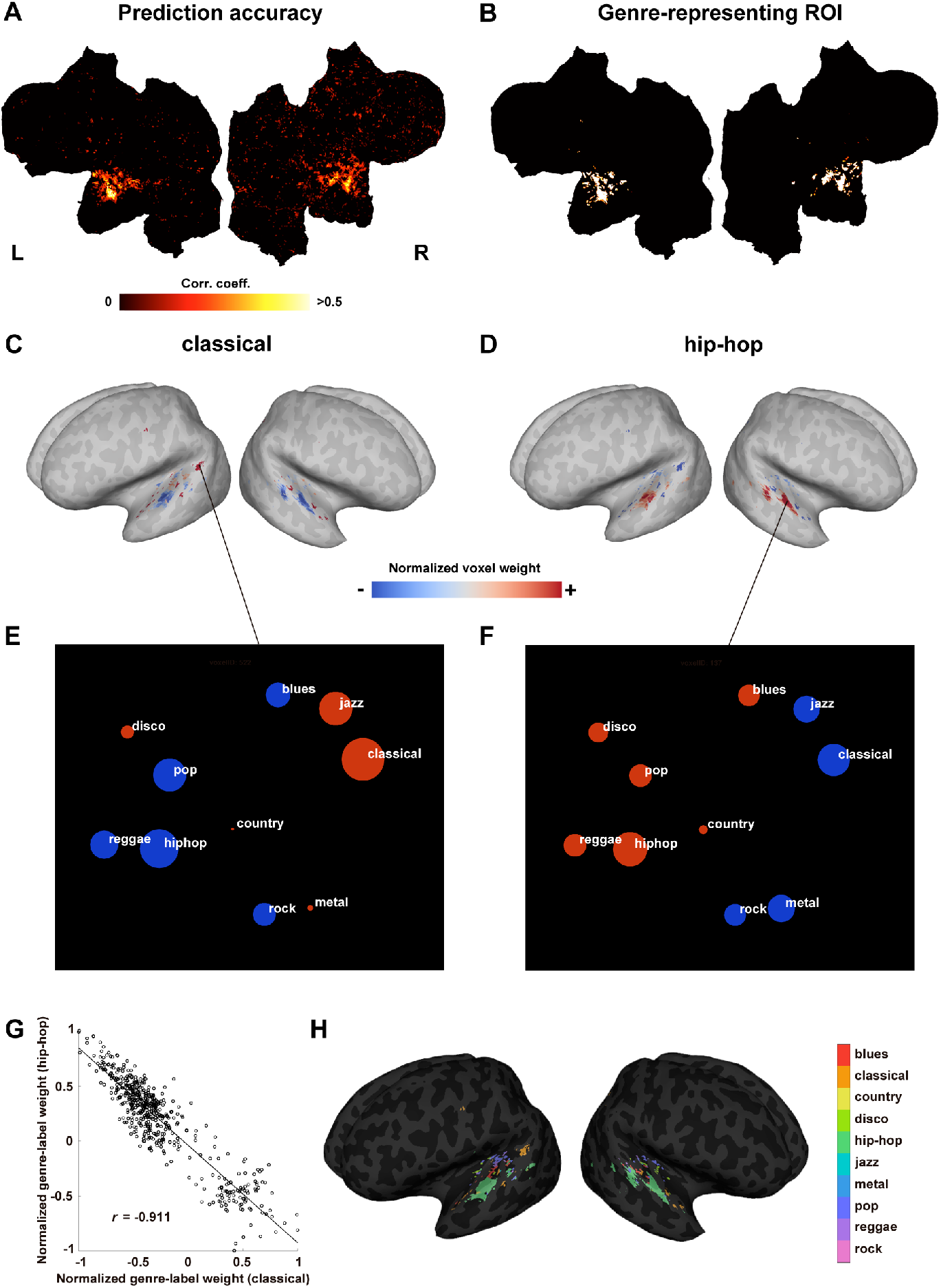
Cortical organization of music genre representations. (A) The cortical map of prediction accuracy using the genre-label model (*p* < 0.05, FDR corrected) is shown on flattened cortical sheets. **(B)** Genre-representing ROI for subject ID01, obtained using the genre-label model. L, Left hemisphere. R, Right hemisphere. **(C,D)** The normalized weights of the genre-label model were projected on the inflated cortical map (genre-weight map; red, positive weight; blue, negative weight), for classical **(C)** and hip-hop music **(D). (E,F)** Weight values for the 10 genres extracted in 2 representative voxels in subject ID01, plotted according to *t*-SNE coordinates. The radius of each circle is equivalent to the weight value of the corresponding music genre at the target voxel (red, positive weight; blue, negative weight). The distance between each circle reflects the differences in cortical representations. **(G)**, Scatterplot showing a negative correlation between the normalized weights in classical and hip-hop music. **(H)**, Cortical map of all music genres tested in the current study for subject ID01. All voxels were assigned 1 of 10 colors according to music genre with the largest weight value in the genre-label model. (See Fig. S3 for maps in other subjects).

To further understand how music genres were represented within the genre-representing ROI, we examined the cortical weight map for each music genre (genre-weight map). Genre-specific cortical organization was observed such that regions positively activated by classical music were negatively activated by hip-hop music (Fig. 2C, D). More quantitatively, the weight values of the genre-label model for classical and hip-hop music showed a clear negative correlation (*p* < 0.001; see Fig. 2G for subject ID01). This opposite representation pattern was consistent across subjects (mean ± sd, *r* = −0.823 ± 0.096).

To illustrate the relative relationships among cortical activations for different music genres, we visualized the weight values of 10 music genres in each voxel using 2-D coordinates derived from *t*-SNE (20). As shown in Fig. 2E, F, we selected the two representative voxels that had the largest weight values for classical (Fig. 2E, voxel ID 522) and hip-hop (Fig. 2F, voxel ID 137) music in subject ID01. This result suggested that classical, jazz, and blues music, or rock and metal music, induced relatively similar activation patterns, whereas classical and hip-hop music seemed to have distinct activation patterns. To investigate the relative contribution of each cortical voxel to the 10 music genres, we further mapped the music genres for which each cortical voxel had the largest weight value (Fig. 2H and Supplementary Fig. 3), which revealed various genre-specific representations within the bilateral STG.

### Genre-specific brain activity was explained by spectro-temporal modulation of music genres

We hypothesized that genre-specific cortical organization in the bilateral STG was produced by the spectro-temporal modulation property of music stimuli. To test this prediction, we first extracted the spectro-temporal modulation energy of each music stimulus using the MTF model (see Methods). By averaging the MTF model features for 48 clips of the same music genre in the training dataset, we obtained distinct spectro-temporal modulations for the 10 music genres (Fig. 3A and Supplementary Fig. 4). For example, classical and hip-hop music showed a clear contrast of opposite spectro-temporal modulation, which is consistent with the distinct activation patterns observed in the genre-label model (Fig. 2C, D, H).

**Fig. 3.**
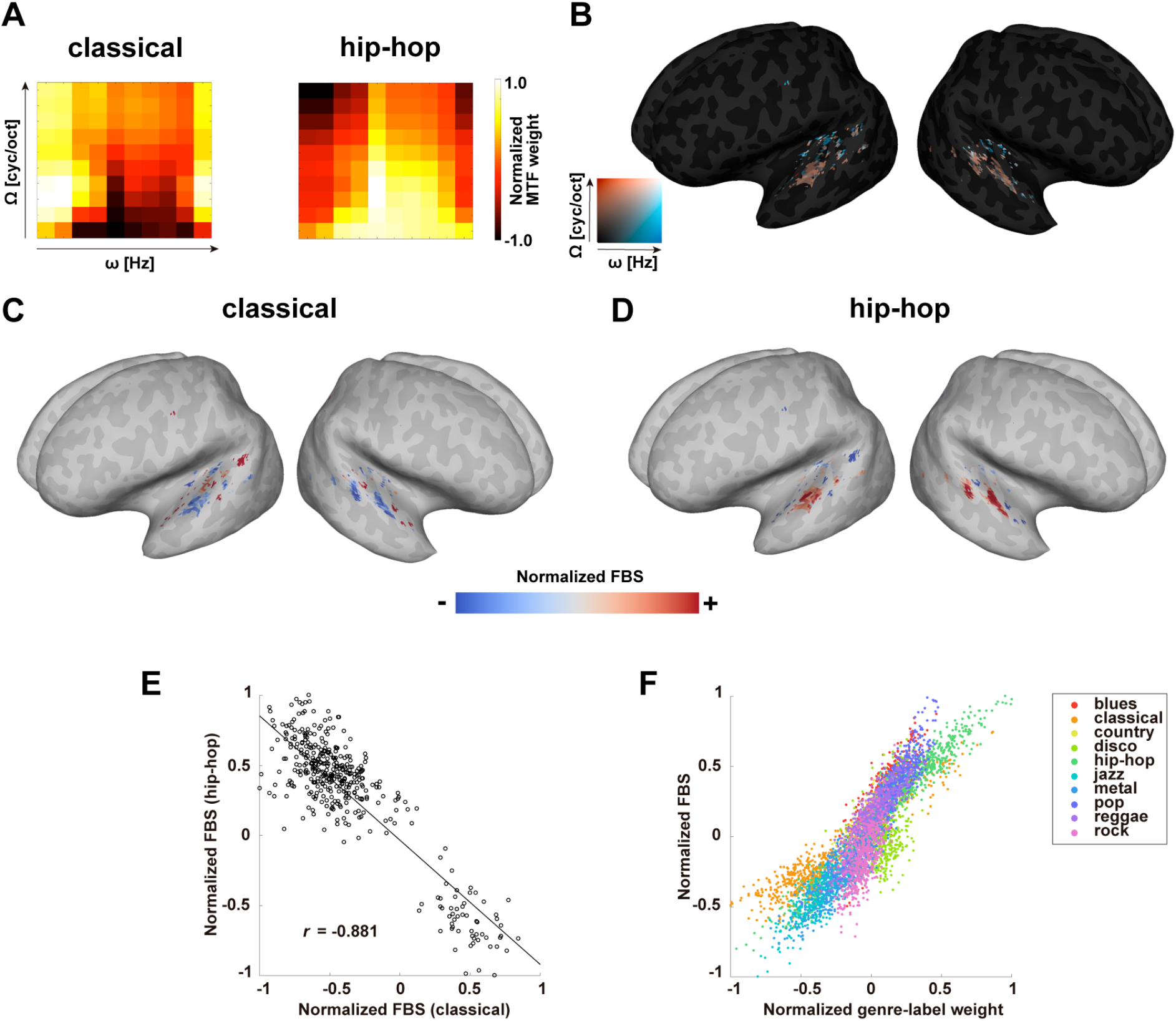
Contribution of spectro-temporal modulations to music genre representations. **(A)** Examples of averaged modulation profiles (classical and hip-hop music). The modulation-transfer function (MTF) model weights were projected on a 2-D plot of the spectral modulation Ω (cyc/oct) and temporal modulation rates ω (Hz). **(B)** Spectro-temporal modulation of cortical voxels in subject ID01. The weight vectors of the MTF model were averaged for 20 central frequencies. Each cortical voxel was assigned the maximum spectral/temporal modulation rate of that voxel. **(C,D)** The feature-brain similarity (FBS) cortical maps of subject ID01 obtained from spectro-temporal modulations of **(C)** classical **(D)** and hip-hop music (i.e., MTF model features). Data are normalized and projected on the inflated cortical map (red, positive weight; blue, negative weight). The genre-wise FBS maps tend to resemble the genre-label weight maps shown in Fig. 2C,D. **(E)** Scatterplot showing the negative correlation between normalized FBS in classical and hip-hop music. **(F)** The 2-D scatterplot of voxel values of the normalized genre-weight map and FBS map of the MTF model (see Fig. 2A,B, and 3C,D, respectively) taken from all voxels in the genre-representing ROI of subject ID01 and overlaid with 10 music genres.

Next, we performed an encoding model fitting with MTF model features. Using the weight matrix of the MTF model, we obtained the spectro-temporal modulation of each cortical voxel (Fig. 3B and Supplementary Fig. 5). By calculating the correlations between spectro-temporal modulations of music genres and those of the cortical voxels, we found a FBS cortical map of each music genre based on its spectro-temporal modulation (Fig. 3C, D). The obtained FBS map reflects the opposite spectro-temporal modulation property of classical and hip-hop music (Fig. 3A), as shown by the negative correlation among FBS of all voxels (*p* < 0.001; see Fig. 3E for subject ID01).

The cortical map obtained in the genre-label model (Fig. 2C,D) and the FBS map based on the MTF model (Fig. 3C,D) were very similar and significantly correlated (Pearson’s correlation coefficient, *p* < 0.001, with Bonferroni correction; Fig. 3F and Supplementary Fig. 6). Significant correlation was consistently observed across all subjects (*r* = 0.826 ± 0.047), indicating that the genre-specific cortical activation patterns were explained by the different spectro-temporal modulations of the 10 music genres.

### Genre classification accuracy based on brain activity, behavior, and MTF model

To investigate any differences in cortical representational specificity among the 10 music genres, we performed a genre classification based on brain activity using a decoding model approach (see Methods). The confusion matrix as well as the classification accuracy (the diagonal elements of the confusion matrix) was evaluated using cortical activation within the genre-representing ROI masks (Fig. 4A and Supplementary Fig. 7). The classification results varied across genres such that classical music was always accurately classified (average classification accuracy, 100%), while performance was poor for rock music across all subjects (16.7 ± 11.8%). We also found that reggae music tended to be classified as hip-hop music (confusion from reggae to hip-hop, 43.3% ± 9.1%), whereas rock music tended to be classified as country music (confusion from rock to country, 36.7% ± 13.9 %).

**Fig. 4.**
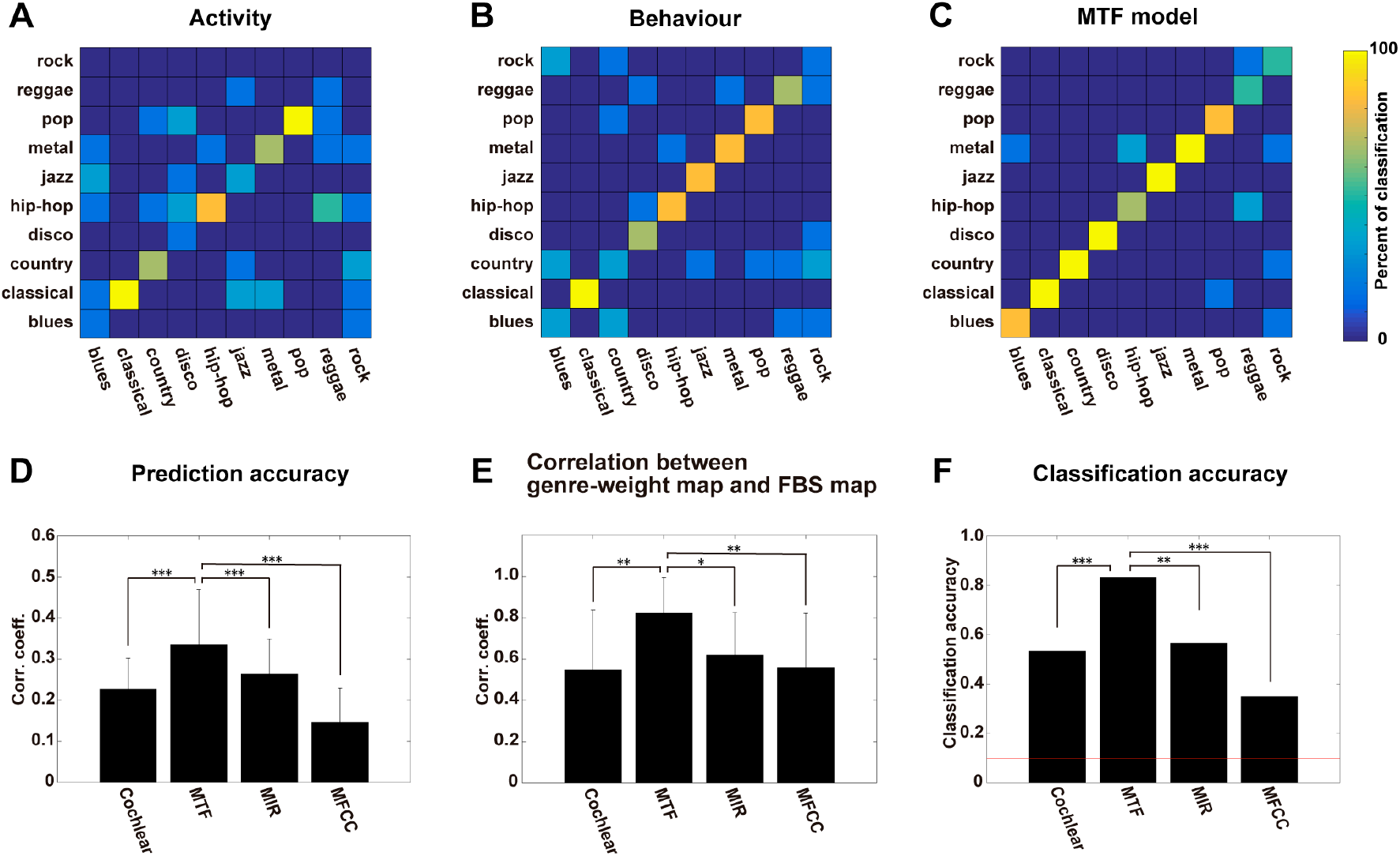
Modulation-transfer function (MTF) model explained genre representational specificities of brain activity and behavior. **(A)** Confusion matrix based on brain activity of subject ID01, using the decoding model approach. For each column of correct music genres, the percentage of classified music clips is plotted on the row of classified music genres. **(B)** Confusion matrix based on behavioral data of subject ID01. **(C)** Confusion matrix based on the MTF model features using the decoding model approach. **(D)** Mean prediction accuracy of the cochlear, MTF, music information retrieval (MIR), and mel-frequency cepstral coefficient (MFCC) models (see Methods for the detailed description of each model), averaged within the genre-representing ROI of subject ID01. **(E)** Pearson’s correlation coefficients between voxels in the genre-weight map and the feature-brain similarity (FBS) map for subject ID01, for the cochlear, MTF, MIR, and MFCC models, averaged for 10 music genres. **(F)** Classification accuracy of the cochlear, MTF, MIR, and MFCC models, with the decoding model approach, averaged for 10 music genres. The red line indicates the chance level (10%). Error bar, sd. * *p* < 0.05, \*\**p* < 0.01, \*\*\**p* < 0.001 (Bonferroni corrected).

To examine how brain activation to music genre was related to behavioral performance during genre classification, we performed an additional behavioral test for each subject who participated in the MRI experiment (MRI subjects) (Fig. 4B and Supplementary Fig. 8). The confusion matrix showed that genre classification performance varied for each genre, such that classical music was always accurately recognized (average classification accuracy, 100%), while rock music was less accurately recognized across all subjects (36.7% ± 13.9%). Behavioral confusion matrices identified brain activity-like error tendencies, such that rock music tended to be classified as country music (confusion from rock to country, 20.0% ± 13.9%). The confusion matrices of behavior and brain activity were significantly correlated for all subjects (Spearman’s correlation coefficient, *ρ* = 0.412 ± 0.049, *p* < 0.001), indicating that genre representational specificity of human behavior mimicked that of brain activity.

As the MRI subjects listened to the music stimuli twice (once in the MRI scanner and again in the behavioral test), there may have been a learning effect. To confirm the generalizability of the behavioral results of these subjects, we recruited an additional 21 subjects for the behavioral tests only (non-MRI subjects, Supplementary Fig. 8). The non-MRI subjects showed variable genre classification accuracy (mean ± sd, 56.3% ± 6.1%; max, 68.3%; min, 43.3%), with performances similar to those of the MRI subjects, such that classical music was accurately recognized (100%) while rock music was not (17.5%). The average behavioral confusion matrices of the MRI subjects and non-MRI subjects were significantly correlated (*ρ* = 0.826, *p* < 0.001).

To further investigate whether the variability in activity-/behavior-based classification accuracies (i.e., representational specificity) was based on the extracted MTF model features, we performed a music genre classification based on these features (Fig. 4C). The classification result was relatively high (mean accuracy across 10 music genres, 83.3%) relative to the activity/behavior-based classification (50.3% and 66.3%, respectively; Wilcoxon signed-rank test, *p* = 0.0020); however, there was notable variation across genres, such that classical music was accurately classified (100%) but performance was low for rock music (50%). We also confirmed that the confusion matrix obtained for the MTF model was significantly correlated with the matrices for brain activity (average Spearman’s correlation coefficients of 5 subjects, *ρ* = 0.521 ± 0.047, *p* < 0.001) and behavior (*ρ* = 0.583 ± 0.054, *p* < 0.001).

### Comparison of four acoustic feature models

To further examine if the MTF model—and not another well-established acoustic feature model—could explain brain activity evoked by the music stimuli, the prediction accuracy was evaluated for the four acoustic feature models (cochlear, MTF, MIR, and MFCC) (Fig. 4D and Supplementary Fig. 9). The MTF model significantly outperformed the other models in all subjects (Wilcoxon signed-rank test, *p* < 0.001 for all subjects, with Bonferroni correction), indicating that it most accurately explained music-evoked brain activity.

We also tested whether the MTF model outperformed the other acoustic models in capturing genre-specific cortical organization. The mean correlation between genre-weight maps and FBS maps confirmed that the MTF model most accurately predicted genre-specific cortical activation patterns (Wilcoxon signed-rank test, *p* < 0.010 for all subjects; Fig. 4E and Supplementary Fig. 10), indicating that the MTF model most accurately captured genre-specific cortical organization in the bilateral STG.

Finally, we tested whether the MTF model outperformed the other models in capturing the representational specificity of various music genres. To this end, we compared the music genre classification accuracy among the four models (cochlear, MTF, MIR, and MFCC) (Fig. 4F). Although the cochlear, MIR, and MFCC models also showed significant correlations with the confusion matrices of the behavior-based approach (average Spearman’s correlation coefficients, cochlear, 0.465; MIR, 0.465; MFCC, 0.343; *p* < 0.0060) and brain activity-based approach (cochlear, 0.459; MIR, 0.513; MFCC, 0.342; *p* < 0.0018), the MTF model significantly outperformed the other models in terms of classification accuracy (Chi-squared test, *p* < 0.0015 vs. each of the other three models), which is consistent with the prediction accuracy and correlation analysis results between genre-weight and FBS maps.

## Discussion

Using fMRI, the present study has demonstrated the cortical organization underlying different music genres. Since the genre-label model did not assume any acoustic properties, the genre-weight maps (Fig. 2C,D,H, and Supplementary Fig. S3) reflect music genre information in the most abstract sense. It was therefore important that we obtained similar weight patterns between the genre-label model and the FBS map based on the MTF model. The FBS map shows how the spectro-temporal modulation of each cortical voxel corresponds to the reference spectro-temporal modulation profile of each music genre. It is thus likely that the weight values in the bilateral STG in the genre-label model were determined by how much each STG voxel’s spectro-temporal modulation property resembles that of the music stimuli.

Previous studies have reported a frequency-selective (i.e., tonotopic) gradient in the STG in humans (21–23). The cochlear model features correspond to the positions in the frequency axis, and may thus reflect the tonotopic property. The MTF model further captures the modulation property around each position in the frequency axis. Spectro-temporal modulation tuning has also been reported in the STG (6, 11–13). Santoro et al. (2014) demonstrated higher performance of the MTF model than the cochlear model for predicting STG activation to natural sound stimuli, which is consistent with the present findings (6). In sum, the spectro-temporal modulations obtained in our study seem to reflect the general processing property of auditory stimuli in the bilateral STG.

Several studies have reported that perceived music genres can be decoded from brain activity. Ghaemmaghami and Sebe (2017) used magnetoencephalogram and electroencephalogram datasets to classify musical stimuli as either pop or rock, using support vector machines (SVM) (24). Casey (2017) and Sengupta et al. (2018) used fMRI data with 5 distinct music genres, then performed activity-based multi-class classification using SVM (25, 26). These studies, however, did not provide how cortical representations of music genres contributed to genre classification. The current results demonstrated underlying mechanisms of such activity-based genre classification.

Music genre classification accuracy was much higher in the MTF model-based approach (83.3%), compared to the behavior (66.3%) or brain activity-based approaches (50.3%). The behavioral results in this study were in the range of previous classification reports (55%–76%) (27, 28) and classification accuracy based on the MTF model was comparable to top-level performances in previous studies using the GTZAN music genre database (29). Moreover, we demonstrated that the confusion matrix of the MTF model was significantly correlated with the matrices of behavior and brain activity (Fig. 4), which indicates that, although the model-based approach was more accurate, human behavior/brain activity and spectro-temporal modulation features show similar classification patterns.

Classification accuracy was examined using four feature models. The MFCC and MIR models were developed in the field of computational science and have been used in previous studies of music-induced brain activity (2, 3, 30). The cochlear model has been used to test cortical activation in the spectral domain (17), although this model cannot capture the dynamic temporal modulation of spectra (see Fig 1C). The MTF model was constructed based on the physiological properties of neurons in the auditory cortex (31) and is widely used in neuroscience research on auditory perception (5–8). Therefore, it is likely that the MTF model is more biologically plausible for addressing the auditory processing of music genres. The results of this study are consistent with this view, as the MTF model exhibited the highest prediction accuracy among the encoding models of cortical activation (Fig. 4D), as well as the highest music genre classification accuracy among the four models on music genre recognition (Fig. 4F).

In the MRI experiments performed in this study, the subjects passively listened to music stimuli and did not perform any genre classification tasks during scanning. It could be argued that we did not confirm that the subjects attentively listened to the stimuli and that we thus overlooked brain regions activated for top-down decisionmaking on music genre classification. To address this concern, we performed behavioral experiments of genre classification for the MRI subjects (Fig. 4A,C) and confirmed that the confusion matrices based on brain activation and behavior were significantly correlated. This result indicates that passive listening to music stimuli captured sufficient brain information for use in behavioral music genre classification.

Some amount of genre-specific organization could be explained by linguistic factors, e.g., most classical and jazz pieces used in the present study were instrumental (i.e., without human voice), while other genres contained the human voice. We have, however, shown that classical and jazz music were not confused with each other (see the confusion matrices in Fig. 4), and that some music genres with voice were not confused with each other (e.g., hip-hop and country music). Moreover, we showed that jazz and blues music were closely represented (Fig. 2E,F), even though jazz was mostly instrumental and blues typically contained voice. Therefore, the effects of linguistic factors cannot solely explain the activation patterns in the bilateral STG

Lastly, we evaluated only handcrafted models (cochlear, MFT, MIR, and MFCC) and did not test the state-of-the-art deep neural network (DNN) model. A recent fMRI study by Kell et al. (2018) that trained a DNN model using a music genre classification task reported that the model could predict cortical activation induced by various sound stimuli (32). However, the authors noted that the DNN model has a disadvantage of interpretability in each unit. As the aim of the present study was to reveal the spectro-temporal modulation basis of music genre representation in the human brain, it seemed reasonable to use handcrafted models.

We thus conclude that music genres are represented in the bilateral STG in a genre-specific way and that representations can be modeled by spectro-temporal modulation profiles extracted from music pieces. The current results indicate the possibility of modeling our abstract categorization of complex auditory stimuli based on the brain activity.

## Methods

### Subjects

Five healthy subjects (referred to as ID01-05; age range 23–33; 2 females; music experience of 4–15 years) with normal hearing participated in the MRI and behavioral experiments. An additional 21 subjects (age 20–24; 5 females; music experience of 0–15 years) participated only in the behavioral experiment. A questionnaire was used to assess subjects’ years of training on their primary instrument, which was used as the index of musical experience. Informed consent was obtained from all subjects prior to their participation. This experiment was approved by the ethics and safety committee of the National Institute of Information and Communications Technology in Osaka, Japan.

### Stimuli and task

Music stimuli from 10 genres (blues, classical, country, disco, hip-hop, jazz, metal, pop, reggae, rock) were randomly taken from the GTZAN music genre dataset (http://marsyasweb.appspot.com/download/data_sets/) (33). For each genre, 54 music pieces (30 s, 22,050 Hz) were selected, resulting in a total of 540 music pieces. From each music piece, a 15-s music clip was randomly selected. For each clip, we applied 2 s of fade-in and fade-out effects and normalized the overall signal intensity in terms of the root mean square (RMS).

Each experiment comprised 18 runs; 12 were considered training runs and 6 were considered test runs. Each run consisted of 40 music clips and had a total duration of 10 min. At the beginning of each run, 15 s of dummy scanning was acquired, which were omitted from analysis. Of the 540 music clips, 480 were used in the training runs and the remaining 60 were reserved for the test runs. In each test run, a set of 10 music clips was presented 4 times in the same order. The clip order was randomized across the experiment. During scanning, subjects fixated on a fixation cross at the center of the screen and listened to the music clips through MRI-compatible earbuds. The experiment was executed over the course of three days, with six runs performed each day.

### MRI data acquisition

Scanning was performed using a 3.0 T MRI scanner (TIM Trio; Siemens, Erlangen, Germany), with a 32-channel head coil. We scanned 68 interleaved axial slices with a thickness of 2.0 mm without a gap using a T2*-weighted gradient echo multi-band echo-planar imaging (MB-EPI) sequence (34) (repetition time (TR) = 1500 ms, echo time (TE) = 30 ms, flip angle (FA) = 62°, field of view (FOV) = 192 × 192 mm^2^, voxel size = 2 × 2 × 2 mm^3^, multiband factor = 4). We obtained 410 volumes in each run. For the anatomical reference, high-resolution T1-weighted images of the whole brain were acquired from all subjects with a magnetization prepared rapid acquisition gradient echo sequence (MPRAGE, TR = 2530 ms, TE = 3.26 ms, FA = 9°, FOV = 256 × 256 mm^2^, voxel size = 1 × 1 × 1 mm^3^).

### Model setting Genre-label model

The genre-label model comprised 10 features corresponding to the 10 music genres. A value of 1 or 0 was assigned to each time bin of 1.5 s to represent whether a music clip of a certain genre was presented.

### Cochlear and MTF models

A sound cochleogram was generated using a bank of 128 overlapping bandpass filters, spanning from 20 to 10,000 Hz (35) (Fig. 1C). The window size was set to 25 ms and the hop size to 10 ms. The filter output averaged across 1.5 s (TR) was used as a feature in the cochlear model.

Next, we extracted features from the MTF model, as done by Chi et al. (2005) (31) (Fig. 1E,F). For each cochleogram, a convolution with modulation-selective filters was calculated. The outputs of two filters with orthogonal phases (quadrature pairs) were squared and summed to yield the local modulation energy (15). The local modulation energy was log-transformed, averaged across 1.5 s, and further averaged within each of the 20 non-overlapping frequency ranges logarithmically spaced in the frequency axis. The filter outputs of the upward and downward sweep directions were averaged. Modulation-selective filters were tuned to 10 spectral modulation scales (Ω = (0.35, 0.50, 0.71, 1.0, 1.41, 2.0, 2.83, 4.0, 5.66, 8.0) cyc/oct) and 10 temporal modulation rates (ω = (2.8, 4.0, 5.7, 8.0, 11.3, 16.0, 22.6, 32.0, 45.3, 64.0) Hz). To reduce the computational burden, the resultant 20 × 10 × 10 = 2000 features were reduced to 302 features using a principal component analysis (PCA), preserving 99% of the variance in the original features.

### MIR and MFCC models

The MIR toolbox was used to extract multiple music-related features from the dataset (36). Consistent with a previous neuroimaging study on music (2), we extracted the following 24 features: RMS energy as the loudness feature; zero-crossing rate, high energy-low energy ratio, spectral centroid, spectral roll-off, spectral entropy, spectral flatness, roughness, spectral spread, spectral flux, and sub-band flux (with nine sub-bands) as timbral features; pulse clarity, fluctuation centroid, and fluctuation entropy as rhythm features; and mode and key clarity as tonal features. For the loudness and timbral features, the frame duration was set to 25 ms, with a 50% overlap between two adjacent frames. For the rhythm and tonal features, the frame duration was set to 3 s with a 33% overlap. Each feature was averaged across 1.5 s. We also used the MIR toolbox to extract MFCC features with 12 channels (36).

### Data analyses

#### fMRI data preprocessing

Motion correction was performed for each run using the Statistical Parametric Mapping toolbox (SPM8; Wellcome Trust Centre for Neuroimaging, London, UK; http://www.fil.ion.ucl.ac.uk/spm/). All volumes were aligned to the first EPI image for each subject. Low-frequency drift was removed using a median filter with a 240-s window. The response for each voxel was normalized by subtracting the mean response and then scaling it to the unit variance. We used FreeSurfer (37, 38) to identify cortical surfaces from anatomical data and to register them to the voxels of the functional data. For each subject, voxels identified in the cerebral cortex were used in the analysis.

#### Voxel-wise encoding model fitting

For each of the above models, cortical activation in each voxel was fitted using a set of linear temporal filters that captured the slow hemodynamic response and its coupling with brain activity (15). A feature matrix F_E_ [T × 4N] was modeled using concatenating sets of [T × N] feature matrices with temporal delays of 1.5, 3, 4.5, and 6 s (T, # of samples; N, # of features). The cortical response R_E_ [T × V] was modeled using the feature matrix F time the weight matrix W_E_ [4N × V] (V, # of voxels):

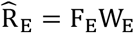

An L2-regularized linear regression using the training dataset (4800 samples, 7200 s) was performed to obtain the weight matrix W_E_. The optimal regularization parameter was evaluated using random resampling of the training dataset into two subsets: 80% of the dataset was used for model fitting and the remaining 20% was used for model validation. This random resampling procedure was repeated 10 times.

The test dataset consisted of 600 samples (900 s). Four repetitions of test datasets were averaged to increase the signal-to-noise ratio. Prediction accuracy was calculated using Pearson’s correlation coefficient between the predicted signal and measured signal in the test dataset. The statistical significance was computed based on the null distribution of correlations between two independent Gaussian random vectors of the same length. The resulting *p* values were corrected for multiple comparisons within each subject using the false discovery rate (FDR) procedure (39). The mean prediction accuracy of each encoding model was calculated by averaging the prediction accuracy of all voxels within the subject-specific region-of-interest mask (see below). All model fitting and analyses were performed using custom software written on MATLAB. For the data visualization on cortical maps, we used pycortex (40).

#### Genre-representing region-of-interest (ROI) mask

We divided the training dataset into training samples (80%) and validation samples (20%). Using the optimal regularization parameter estimated in the analysis of the genre-label model, we repeated the model fitting 50 times. Voxels showing significant prediction accuracy (FDR corrected) for more than 80% of repetitions were selected for the ROI mask.

#### Visualization of relative relationships among music genres

To visualize the relative relationships among music genres according to cortical representations, we used *t*-distributed stochastic neighbor embedding (*t*-SNE) (20). Specifically, for each voxel within the ROI mask, we extracted the estimated weight vectors of the genre-label encoding models. To obtain a general result across participants, we concatenated weight vectors of five participants. We then performed dimensionality reduction on the aggregated weight vectors, using *t*-SNE to obtain the relative locations of 10 music genres in a 2-D space (perplexity = 5).

#### Decoding of genre-labels

In the decoding model, the cortical response matrix RD [T × 4V] was modeled using concatenating sets of [T × V] matrices with temporal delays of 1.5, 3, 4.5, and 6 s. The genre-label matrix G [T × 10] was modeled using the neural response matrix R_D_ times the weight matrix W_D_ [4V × 10]:

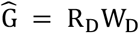

The weight matrix WD was estimated using an L2-regularized linear regression with the training dataset following the same procedure used for the encoding model fitting. To calculate the classification accuracy, we first assigned genre-label indices (1 to 10) to each time point by taking the argmax of the decoded genre-label matrix. The representative genre-label index for each music clip was estimated using the majority voting method (41).

In the activity-based approach, we obtained a response matrix RD for each subject as well as for the combined data of all subjects. The combined response matrix was obtained by concatenating the cortical activation of three subjects within the subject-specific ROI masks. In the feature based approach, the feature matrix F_D_ [T × N] was used instead of the response matrix R_D_.

#### Spectro-temporal modulation of music genres

Spectro-temporal modulation of each music genre was assessed based on the feature matrix F_E_ used in the encoding model fitting. The MTF model feature matrix was restored to the original size (F_E_U^t^) by multiplying it by the transposed PCA coefficient matrix U^t^. The MTF model weight matrix W_E_ was also transformed by multiplying it by the PCA coefficient matrix U. The genre-specific feature vector was calculated by averaging the MTF model features of the 48 clips in the training dataset for each genre. For visualization of the MTF model, we further averaged the feature values obtained at 20 central frequencies for each of the 10 × 10 combinations of spectral modulation Ω (cyc/oct) and temporal modulation ω (Hz). The genre-specific feature vector was calculated for the other models using the same procedure.

#### Spectro-temporal modulation of cortical voxels

Spectro-temporal modulation of each cortical voxel was assessed using the transformed weight matrix UW_E_. The voxel-specific weight vector was calculated by averaging the MTF model weight values of the 48 training clips for each genre. For visualization of the MTF model, we further averaged the weight values obtained at 20 central frequencies for each of the 10 × 10 combinations of spectral modulation Ω (cyc/oct) and temporal modulation ω (Hz). The voxel-specific weight vector was calculated for the other models using the same procedure.

#### Correlations between spectro-temporal modulations of music genres and cortical voxels

To examine whether genre-representing activation patterns were explained by the extracted features, we calculated the Pearson’s correlation coefficients between spectro-temporal modulations of music genres and cortical voxels. The resultant 10 cortical maps were defined as feature-brain similarity (FBS) maps. We further calculated Pearson’s correlation coefficients between cortical weights in the genre-label model and those in the FBS map for each of the 10 music genres.

### Behavioral experiment

To confirm that brain activation to music genres was related to behavioral performance of genre classification, we performed additional behavioral experiments. Subjects were first asked to listen to 3 original music clips (30 s) per genre, which were randomly selected from the 460 clips not used in the MRI experiment. In the training session, subjects were asked to understand the property of each music genre. Subjects then listened to the 60 music clips used as the validation dataset in the MRI experiment and judged what music genre the target music clip belonged to by filling in 1 of 10 cells on the answer sheet. Subjects listened to each music clip only once and in the same presentation order as in the fMRI experiment. These behavioral experiments were performed in a soundproof room by the same subjects who participated in the MRI experiments, as well as an additional 21 subjects who did not participate in the MRI experiments. Of the 21 non-MRI subjects, data from one subject was excluded because their average accuracy (30.0%) was outside the mean ± 3*sd range (and also outside the median 3*interquartile range) of all subjects.

## Acknowledgments

We are grateful to MEXT/JSPS KAKENHI (Grant Number JP18H05091 in #4903 (Evolinguistics), JP15H05311) for the partial financial support of this study. The funders had no role in study design, data collection and analysis, decision to publish, or preparation of the manuscript.

## Author contribution

T.N., N.K., and S.N. designed the study; T.N. and N.K. carried out the experiment; T.N. analyzed the data with support from N.K.; T.N., N.K., and S.N. wrote the manuscript.

## Data availability

The data that support the findings of this study are available from the corresponding author upon request.

## References

1. Sturm BL (2012) A Survey of Evaluation in Music Genre Recognition. Adaptive Multimedia Retrieval: Semantics, Context, and Adaptation, Lecture Notes in Computer Science., eds Nürnberger A, Stober S, Larsen B, Detyniecki M (Springer, Cham, Cham), pp 29–66.

2. Alluri V, et al. (2012) Large-scale brain networks emerge from dynamic processing of musical timbre, key and rhythm. Neuroimage 59(4):3677–3689.

3. Toiviainen P, Alluri V, Brattico E, Wallentin M, Vuust P (2014) Capturing the musical brain with Lasso: Dynamic decoding of musical features from fMRI data. Neuroimage 88:170–180.

4. Hoefle S, et al. (2018) Identifying musical pieces from fMRI data using encoding and decoding models. Sci Rep 8(1):2266.

5. Patil K, Pressnitzer D, Shamma S, Elhilali M (2012) Music in Our Ears: The Biological Bases of Musical Timbre Perception. PLoS Comput Biol 8(11):e1002759.

6. Santoro R, et al. (2014) Encoding of Natural Sounds at Multiple Spectral and Temporal Resolutions in the Human Auditory Cortex. PLoS Comput Biol 10(1):e1003412.

7. Norman-Haignere S, Kanwisher NG, McDermott JH (2015) Distinct Cortical Pathways for Music and Speech Revealed by Hypothesis-Free Voxel Decomposition. Neuron 88(6):1281–1296.

8. Santoro R, et al. (2017) Reconstructing the spectrotemporal modulations of real-life sounds from fMRI response patterns. Proc Natl Acad Sci 114(18):4799–4804.

9. Depireux DA, Simon JZ, Klein DJ, Shamma SA (2001) Spectro-Temporal Response Field Characterization With Dynamic Ripples in Ferret Primary Auditory Cortex. J Neurophysiol 85(3):1220–1234.

10. Pasley BN, et al. (2012) Reconstructing speech from human auditory cortex. PLoS Biol 10(1):e1001251.

11. Hullett PW, Hamilton LS, Mesgarani N, Schreiner CE, Chang EF (2016) Human Superior Temporal Gyrus Organization of Spectrotemporal Modulation Tuning Derived from Speech Stimuli. J Neurosci 36(6):2014–2026.

12. Schonwiesner M, Zatorre RJ (2009) Spectro-temporal modulation transfer function of single voxels in the human auditory cortex measured with high-resolution fMRI. Proc Natl Acad Sci 106(34):14611–14616.

13. Langers DRM, Backes WH, Van Dijk P (2003) Spectrotemporal features of the auditory cortex: The activation in response to dynamic ripples. Neuroimage 20(1):265–275.

14. Naselaris T, Kay KN, Nishimoto S, Gallant JL (2011) Encoding and decoding in fMRI. Neuroimage 56(2):400–410.

15. Nishimoto S, et al. (2011) Reconstructing visual experiences from brain activity evoked by natural movies. Curr Biol 21(19):1641–1646.

16. Kay KN, Naselaris T, Prenger RJ, Gallant JL (2008) Identifying natural images from human brain activity. Nature 452(7185):352–355.

17. de Heer WA, Huth AG, Griffiths TL, Gallant JL, Theunissen FE (2017) The Hierarchical Cortical Organization of Human Speech Processing. J Neurosci 37(27):6539–6557.

18. Huth AG, De Heer WA, Griffiths TL, Theunissen FE, Gallant JL (2016) Natural speech reveals the semantic maps that tile human cerebral cortex. Nature 532(7600):453–458.

19. Nakai T, Koide-Majima N, Nishimoto S Encoding and decoding of music-genre representations in the human brain. Proceedings of the IEEE International Conference on Systems, Man, and Cybernetics, p in press.

20. van der Maaten L, Hinton G (2008) Visualizing Data using t-SNE. J Mach Learn Res 9:2579–2605.

21. Moerel M, De Martino F, Formisano E (2014) An anatomical and functional topography of human auditory cortical areas. Front Neurosci 8(8):225.

22. Leaver AM, Rauschecker JP (2016) Functional Topography of Human Auditory Cortex. J Neurosci 36(4):1416–1428.

23. Ahveninen J, et al. (2016) Intracortical depth analyses of frequency-sensitive regions of human auditory cortex using 7TfMRI. Neuroimage 143:116–127.

24. Ghaemmaghami P, Sebe N (2016) Brain and music: Music genre classification using brain signals. EUSIPCO (IEEE), pp 708–712.

25. Sengupta A, Pollmann S, Hanke M (2018) Spatial band-pass filtering aids decoding musical genres from auditory cortex 7T fMRI. F1000Research 7:142.

26. Casey MA (2017) Music of the 7Ts: Predicting and Decoding Multivoxel fMRI Responses with Acoustic, Schematic, and Categorical Music Features. Front Psychol 8:1179.

27. Lippens S, Martens JP, De Mulder T (2004) A comparison of human and automatic musical genre classification. 2004 IEEE International Conference on Acoustics, Speech, and Signal Processing (IEEE), p iv-233–iv-236.

28. Seyerlehner K, Widmer G, Knees P (2011) A Comparison of Human, Automatic and Collaborative Music Genre Classification and User Centric Evaluation of Genre Classification Systems. Adaptive Multimedia Retrieval. Context, Exploration, and Fusion., eds Detyniecki M, Knees P, Nürnberger A, Schedl M, Stober S (Springer Berlin / Heidelberg), pp 118–131.

29. Sturm BL (2013) The GTZAN dataset: Its contents, its faults, their effects on evaluation, and its future use. arXiv preprint arXiv:1306.1461.

30. Güçlü U, Thielen J, Hanke M, van Gerven MAJ Brains on Beats. Advances in Neural Information Processing Systems, pp 2101–2109.

31. Chi T, Ru P, Shamma S a. (2005) Multiresolution spectrotemporal analysis of complex sounds. J Acoust Soc Am 118(2):887.

32. Kell AJE, Yamins DLK, Shook EN, Norman-Haignere S V., McDermott JH (2018) A Task-Optimized Neural Network Replicates Human Auditory Behavior, Predicts Brain Responses, and Reveals a Cortical Processing Hierarchy. Neuron 98(3):630–644.

33. Tzanetakis G, Cook P (2002) Musical genre classification of audio signals. IEEE Trans Speech Audio Process 10(5):293–302.

34. Moeller S, et al. (2010) Multiband multislice GE-EPI at 7 tesla, with 16-fold acceleration using partial parallel imaging with application to high spatial and temporal whole-brain FMRI. Magn Reson Med 63(5):1144–1153.

35. Ellis DPW (2009) Gammatone-like spectrograms. web Resour http://www.ee.columbia.edu/~dpwe/resources/matlab/.

36. Lartillot O, Toiviainen P, Eerola T (2008) A Matlab Toolbox for Music Information Retrieval. Data Analysis, Machine Learning and Applications (Springer, Berlin, Heidelberg.), pp 261–268.

37. Dale AM, Fischl B, Sereno MI (1999) Cortical surface-based analysis: I. Segmentation and surface reconstruction. Neuroimage 9(2):179–194.

38. Fischl B, Sereno MI, Dale AM (1999) Cortical Surface-Based Analysis II: inflation, flattening, and a surface-based coordinate system. Neuroimage 9(2):195–207.

39. Benjamini Y, Hochberg Y (1995) Controlling the false discovery rate: a practical and powerful approach to multiple testing. J R Stat Soc B 57(1):289–300.

40. Gao JS, Huth AG, Lescroart MD, Gallant JL (2015) Pycortex: an interactive surface visualizer for fMRI. Front Neuroinform 9. doi:10.3389/fninf.2015.00023.

41. Dalwon J, Minho J, Yoo CD (2008) Music genre classification using novel features and a weighted voting method. 2008 IEEE International Conference on Multimedia and Expo (IEEE), pp 1377–1380.

